# Near-Physiological in vitro Assembly of 50S Ribosomes Involves Parallel Pathways

**DOI:** 10.1101/2022.11.09.515862

**Authors:** Xiyu Dong, Lili K. Doerfel, Kai Sheng, Jessica N. Rabuck-Gibbons, Anna M. Popova, Dmitry Lyumkis, James R. Williamson

## Abstract

Understanding the assembly principles of biological macromolecular complexes remains a significant challenge, due to the complexity of the systems and the difficulties in developing experimental approaches. As a ribonucleoprotein complex, the ribosome serves as an ideal model system for the profiling of macromolecular complex assembly. In this work, we report an ensemble of large ribosomal subunit intermediate structures that accumulate during synthesis in a near-physiological and co-transcriptional *in vitro* reconstitution system. Thirteen pre-50S intermediate maps covering the whole assembly process were resolved using cryo-EM single particle analysis and heterogeneous subclassification. Segmentation of the set of density maps reveals that the 50S ribosome intermediates assemble based on fourteen cooperative assembly blocks, including the smallest assembly core reported up to now, which is composed of a 600-nucleotide-long folded rRNA and three ribosomal proteins. The cooperative blocks assemble onto the assembly core following a defined set of dependencies, revealing the parallel assembly pathways at both early and late assembly stages of the 50S subunit.

## INTRODUCTION

The ribosome is the largest and one of the most complicated enzymes in cells, making up 25% of the cell’s dry mass (1), and is responsible for protein synthesis in all living organisms on earth. The bacterial ribosome has a total mass of over 2.5 MDa and is composed of 3 ribosomal RNAs and 54 ribosomal proteins. The process by which ribosomal components interact with each other and assemble into the functional ribosomes is termed ribosome biogenesis. During bacterial ribosome biogenesis, the ∼5 kb primary rRNA transcript is cleaved into the three ribosomal RNAs, undergoes post-transcriptional chemical modifications by a set of enzymes, folds into an intricate secondary and tertiary structure, and binds to the ribosomal proteins, all facilitated by dozens of ribosome biogenesis factors(2, 3). Despite its complexity, the bacterial ribosome assembly process takes approximately 2 minutes in cells(4, 5), so the intermediates during the step-wise ribosome assembly process are incredibly short-lived, and intermediates are present at ∼2% of the entire ribosome population in rapidly growing bacterial cells (6), making it difficult to directly separate and investigate the incomplete ribosomes under native physiological environments.

To characterize the incomplete ribosome assembly intermediates, genetic (7-11) and biochemical perturbations (12), in combination with newly developed structural tools, have been employed to provide insights into the features of the intermediates and putative assembly pathways. A series of intermediates resulting from bL17-depletion strain were previously described, (13), which revealed five cooperative assembly blocks, and evidence for re-routed assembly pathways under the perturbation of ribosomal protein depletion. Thus far, the studies of the LSU have mostly focused on the late assembly stage (7-11, 14, 15), making the early assembly of LSU a remaining mystery.

The classical *in vitro* reconstitution of 50S subunit was first achieved by Nierhaus and Dohme (16). Recently, a set of five structures assembled using this protocol provided insights into the final maturation of the peptidyl transferase center (17). While this remarkable *in vitro* reconstitution has provided many intriguing insights into 50S assembly (17-19), the reconstitution reaction is carried out with fully processed and modified RNAs under non-physiological buffer and temperature conditions. Further, reconstitution is carried out in the absence of assembly factors, making it unsuitable for analysis of the role of specific factors.

In 2013, Jewett et al. reported an *in vitro* system named integrated rRNA synthesis, ribosome assembly, and translation (iSAT), which provides a defined system for ribosome biogenesis in a near-physiological environment (20-23). The reagents added into an iSAT reaction include the plasmid encoding the rRNA operon and a reporter gene, the cell extract that contains factors and chaperones needed for ribosome biogenesis, and the complete set of purified ribosomal proteins. By monitoring the production of a reporter protein, such as GFP or luciferase, the translation process can be observed in real time. In comparison with the classical *in vitro* reconstitution protocol, iSAT is carried out at near-physiological environment (37 °C, 10 mM Mg^2+^) and allows the cascade of transcription, ribosome assembly, and translation to be monitored directly. Importantly, the ribosomes assembled in iSAT are active without the need for additional treatments.

The iSAT reaction presents a powerful opportunity to study ribosome biogenesis due to the ability to conveniently control every component involved in the ribosome assembly process without the concern of its lethality to the organism. In addition, the course of the assembly process can be monitored over time. In this work, we succeeded in harnessing iSAT to capture the very early intermediates during assembly of ribosomal large subunit and solved their structures by Cryo-EM. The set of intermediates can be arranged into a putative temporal order, spanning assembly stages from the earliest folding of the 5’-terminal domain I to later stages of nearly completed subunits. The landscape exhibits both parallel and sequential folding of cooperative domains, providing a robust process to ensure efficient assembly.

## MATERIAL AND METHODS

### Purification of native ribosomes and S150 extract

Native ribosomes and S150 extract were purified from S30 crude cell extract as described (21). Briefly, *E. coli* MRE cells were lysed and S30 crude extract was obtained by two clarification spins. The S30 crude extract was layered onto a 10-40% sucrose cushion and ultracentrifugation at 90,000 g and 4°C for 20 h, resulting in a ribosome containing pellet and S150 containing supernatant. Ribosomes were resolved in buffer C (10 mM TrisOAc (pH 7.5 at 4°C), 60 mM NH_4_Cl, 7.5 mM Mg(OAc)_2_, 0.5 mM EDTA, 2 mM DTT) and stored at -80°C. The concentration of 70S ribosomes was determined by A260 readings (1 A260 unit = 24 pmol 70S) (24).

The supernatant was centrifuged at 150,000 g and 4°C for 3 h and the top two-thirds of the supernatant was recovered. Using Snake Skin Dialysis tubing (3500 Da MWCO, Thermo fisher) The S150 extract was dialyzed into the AEB storage buffer (10 mM TrisOAc, pH 7.5 at 4°C, 20 mM NH_4_OAc, 30 mM KOAc, 200 mM KGlu,10 mM Mg(OAc)_2_, 1 mM Spermidine, 1 mM Putrescine, 1 mM DTT) and concentrated using a 3 kD MWCO centricon (Merck Millipore). For best performance in the iSAT reaction, the A260 and A280 of the S150 extract was targeted to be 25 OD and 15 OD, respectively. The protein concentration in the S150 extract was determined by Bradford assay (Thermo Fischer scientific). The S150 extract was stored in small aliquots at - 80°C.

### Purification of ribosomal proteins

Ribosomal proteins were purified using existing protocols (21, 24) with slight modifications: 70S ribosomes were mixed with an equal volume of buffer M (25 mM Tris-HCl pH 7.5, 20 mM MgCl_2_, 100 mM KCl, 2 mM DTT) containing 8 M Urea and 6 M LiCl, and incubated on ice overnight. After centrifugation at 16,000 g for 15 min at 4 °C, the supernatant, containing ribosomal proteins, was collected. The pellet was washed with buffer M containing 8 M Urea and 6 M LiCl, incubated on ice for at least 1hour, and after centrifugation at 16,000 g for 15 min at 4 °C, the supernatant was collected. The combined supernatants were dialyzed twice against 100 volumes of buffer M containing 1 M KCl (for 6 hours and overnight) using a 1000 DaMWCO Tube-O-Dialyzers (G-Biosciences). The concentration of TP 70 was determined by measuring A230 (1unit A230 = 240pmol TP70). Aliquots were flash frozen and stored at -80°C.

### iSAT reaction

The iSAT reaction was performed as described previously (20, 21) with slight modifications. Briefly, the iSAT reaction was performed in 57 mM Hepes-KOH, 1.5 mM Spermidine, 1 mM Putrescine, 10 mM Mg(Glu)_2_, and 150 mM KGlu at pH 7.5 with 2 mM DTT, 0.33 mM NAD, 0.27 mM CoA, 4 mM Oxalic acid, 2% (w/v) PEG-6000, 2 mM amino acids (Roche), 1 nM pY71sfGFP plasmid encoding superfolder GFP (M. Jewett), 0.1mg/mL T7 RNA polymerase, 42 mM phosphoenolpyruvate (Roche) and NTP+ mix (1.6 mM ATP (Sigma), 1.15 mM of GTP, CTP and UTP each (Sigma), 45.3 μg/μL tRNA from E. coli MRE 600 (Roche), 227.5 μg/μL Folinic acid pH 7.2). The above components were premixed. The final concentration with respect to the total volume of the iSAT reaction is provided. A 5.6µL aliquot of the premix was pipetted into 6 µl of S150 extract. Ribosomal proteins and plasmid encoding the rrnB operon (pT7rrnb; provided by M. Jewett) were added to a final concentration of 0.4 µM and 4 nM respectively. iSAT reactions of 15 μL each were performed in 96 well plates (Applied Biosystems) and incubated in a StepOnePlus Real-time PCR System (Applied Biosystems) at 37 °C for variable time. sfGFP production was detected by fluorescence measurement at 5 min intervals (excitation: 450–490 nm, emission: 510– 530 nm).

### Electron microscopy sample preparation

iSAT reactions were diluted approximately three-fold with buffer E (50 mM Tris pH 7.8, 10 mM MgCl_2_, 100 mM NH_4_Cl, 6 mM β-mercaptoethanol) and loaded onto a 10-40% (w/v) sucrose gradient in buffer E. Gradients were spun in a Beckman SW41 rotor at 26,000 rpm for 12 h and 4°C. Gradient fractions containing ribosomal particles as indicated by A260 readings and Agarose gel were spin-concentrated using a 30 kDa MW cutoff filter (Amicon) and the buffer was exchanged to buffer A (20 mM Tris-HCl, 100 mM NH_4_Cl, 10 mM MgCl_2_, 0.5 mM EDTA, 6 mM β-mercaptoethanol; pH 7.5). 3 uL sample was applied to a plasma cleaned (Gatan, Solarus) 1.2 mm hole, 1.3 mm spacing holey gold grid and manually plunge frozen in liquid ethane (25).

### Electron microscopy data collection

Single particle data were collected using Leginon software on a Titan Krios electron microscope (FEI) operating at 300 keV equipped with a K2 Summit direct detector (Gatan) with a pixel size of 1.31 Å at 22,500 magnification. A dose of rate of ∼5.8 e^-^/pix/sec was collected across 60 frames with a dose of ∼50 e^-^/Å^2^. To overcome problems of preferred orientation, the data was collected at the tilt of -20° (26). A total of 4607 micrographs (15 min: 1655; 35min: 837; 71 min: 828; 130 min: 565; 240 min: 722) were collected.

### Electron microscopy data processing

The micrograph frames were aligned using MotionCor2 (27) in the Appion image processing wrapper (28). The subsequent data processing was all performed in cryoSPARC (29). The micrographs from five iSAT time course datasets were imported into cryoSPARC separately. The CTF parameters were estimated using CTFFIND4 (30). A total of 1,120,554 particles were automatically picked and extracted from five datasets with default parameters. Two rounds of 2D classification were performed, where 30S and 70S particles are removed from the dataset. 900,883 particles were selected and used for the iterative subclassifications.

For ribosome assembly intermediate datasets, the goal of data processing is to reconstruct a diverse set of as many intermediate classes as possible at an interpretable resolution. An iterative subclassification approach (31) was used where ab initio reconstructions were performed to reach the above goal. The first round of ab initio reconstructions was performed for 5 classes. The classes displaying clear RNA structural features were selected for the subsequent subclassification and reconstruction. The remaining classes were subjected to another round of ab initio reconstruction to confirm that the respective particles could be excluded from further analysis. The selected classes were then sub-classified by iterative ab initio reconstructions. The number of subclasses specified at any given stage depend on the number of the particles in the class (more than 100 k: 5 classes, 100 k-10 k: 4 classes, less than 10 k: 3 classes). The hierarchical connections between the ab initio reconstruction jobs during the iterative subclassification are listed in Table S3. When the number of particles in a reconstructed class is less than 2000, subclassification was terminated and the particles of the class were subjected to homogeneous refinement. All parameter settings for refinement were default in cryoSPARC. Classes that failed to provide a high-resolution reconstruction (worse than 10 Å) were discarded. After the iterative sub-classification, a total of 102,863 particles were reconstructed into 30 density maps.

During the iterative sub-classification, the major classes are independently divided into smaller classes, and particles that are mis-assigned early in subclassification can be separated out and reconstructed during the later sub-classifications. Therefore, it is possible that similar maps can emerge from different initial major classes, and these similar maps should be identified and combined. To determine the differences between the set of 30 maps, the pairwise molecular weight difference was calculated among all the thirty structures, and these differences were used as the metric for hierarchical cluster analysis, as described before (31). A 10.0 kDa threshold was chosen for identifying maps that were essentially similar, and the particles from such classes were combined, repeating the ab initio reconstruction (with 1 class) and homogenous refinement. After clustering and combining, thirteen distinct density maps were reconstructed with global resolutions ranging from 4.53 Å to 8.83 Å.

### Quantitation of rRNA and protein occupancy in EM density maps

The crystal structure of *E*.*coli* 50S subunit (PDB: 4YBB) was segmented into 139 elements and binarized to serve as a reference surface (13, 31). The density maps of iSAT time course intermediates were binarized with the threshold voxel level = 1. The relative values between the density maps and the reference surface maps were calculated, resulting in the occupancy values between 0 and 1 for each element.

The occupancy value for each element was binarized according to two thresholds calculated from the mean and standard deviation of the occupancy values. For each element, the occupancy values of each class were plotted and sorted by increasing value. The three-step increment of the occupancy values was calculated and the maximum was labeled as the peak (y). The upper threshold (u) is the mean of the occupancy values above the peak, minus the standard deviation of the occupancy above the peak. The lower threshold (l) is the mean of the occupancy values below the peak plus the standard deviation of the occupancy values below the peak. If y ≤ 1, then l = u/2. If u < l, then define l’ = (u + l) / 2 = u’, and use l’ and u’ as the thresholds. If u < 0.4, u = s_max_ / 2, l = u / 2. The occupancy values that are larger than the upper threshold were binarized to 1, while the occupancy values that are smaller than the lower threshold were binarized to 0, others (between the upper threshold and the lower threshold) were determined by manually comparing the electron density of the element among all density maps in Chimera. The original code for binarization is accessible at GitHub.

### Element correlation analysis

The correlation between occupancy for the elements (rRNA helices and ribosomal proteins) was determined pairwise by a quadrant analysis. A scatterplot was generated of the occupancy values for element i and element j of all density maps. As the occupancy values were binarized according to the procedure above, the points on the scatter plot will have four possible coordinate values: (0,0), (0,1), (1,0), (1,1). If the coordinates of the dots are only (0,0) and (1,1), the two elements were always resolved or not resolved together in the same density maps, the two elements are considered to be correlated. All the elements correlated to each other were combined and defined as a cooperative assembly block. The original code for elements correlation analysis is accessible at GitHub.

### Dependency analysis for the cooperative assembly blocks

The quadrant analysis described above was also used to analyze the dependency between the cooperative assembly blocks. The dots of the elements that were defined into a cooperative assembly block could be treated as one point. For each scatter plot, if the coordinates of the points are only (0,0), (0,1), (1,1) or (0,1), (1,1), then block i is considered to be dependent on the presence of block j. If the coordinates of the dots are only (0,0), (1,0), (1,1) or (1,0), (1,1), block j is considered to be dependent on block i.

In the dependency map shown in Figure 4, the arrows indicate the dependencies between the blocks. For a parsimonious and clear view of the dependency map, only the closest dependencies were plotted. For example, if block c has dependencies a → c and b → c, and block b has dependency a → b, the a → c dependency is pruned, giving the concise dependencies a → b → c. The original code for block dependency analysis is accessible at GitHub.

### Quantitative mass spectrometry of ribosomal proteins

To quantify the protein composition of iSAT ribosomes, ribosomal proteins were prepared as described (13). In brief, 50 pmol of ^14^N labelled iSAT ribosomes were mixed with an equal amount of ^15^N labelled WT ribosomes purified from *E. coli* MRE600 cells. Proteins were precipitated in 13 % trichloroacetic acid (TCA) overnight at 4 °C, pelleted by centrifugation at 14,000 rpm for 30 minutes at 4 °C. The pellet was washed once in 700 μL cold 10% TCA and once in 700 μL cold acetone. The air-dried pellet was re-dissolved in 40 μL 100 mM NH_4_HCO_3_, 5% acetonitrile (ACN), 5 mM DTT and incubated at 65 °C for 10 minutes. Subsequently, 10 mM iodoacetamide (IAA) was added and the sample was incubated at 30 °C for 30 minutes in the dark. The proteins were digested by 0.2 μg modified grade porcine trypsin (Promega, Co., Madison, WI) during overnight incubation at 37 °C. The next day, peptides were purified using Pierce C18 Spin column (Thermo scientific) following the manufacturers protocol. Peptides were eluted from the column by adding 60 μL of 70 % ACN, 0.1% formic acid and the eluent was dried in a Speed-Vac concentrator. Prior to MS analysis, the samples were re-dissolved in 10 μL MS Buffer (5% ACN, 0.1% formic acid) and centrifuged for 10min at high speed. 4 μL sample were mixed with 1 pmol of a retention time calibration mixture (IRT, Pierce).

SWATH-MS was acquired on a Sciex 5600+ TripleTOF mass spectrometer coupled to an Eksigent nano-LC 400 system. Using the auto sampler, peptides were injected onto a SB-C18 Nano HPLC column (0.5 × 150 mm, Agilent Zorbax), and resolved using a 120min gradient from 5-45% acetonitrile in 0.1% formic acid. SWATH data-independent acquisition was obtained in positive ion mode over the 400-1200 m/z range. The mass spectral data analysis and r-protein quantification against the 15N-labeled reference was performed using Skyline (32).

### Quantitative mass spectrometry of rRNA modifications

Quantitative analysis of rRNA modifications in iSAT ribosomes has been carried out relative to modifications present in 70S rRNA isolated and purified from ^15^N-labeled *E. coli* MRE600 cells. rRNA isotope labeling, isolation and purification using sucrose gradient ultracentrifugation has been carried out as described in prior work (33). Replicate analysis has been performed using two independent iSAT preparations, from which sucrose fractions corresponding to iSAT 70S ribosomes have been combined and used for LC-MS sample preparation. For Replicate 1, isolated and purified iSAT rRNA were combined with wild-type ^15^N-labeled rRNA in about equal molar ratio based on A260, digested using RNase T1 or A for 1 h at 55 °C in 25 mM ammonium acetate (pH = 6), then subjected to LC-MS analysis. For Replicate 2, iSAT rRNA were first combined with wild-type ^15^N-labeled rRNA and then derivatized with CMCT (N-Cyclohexyl-N′-(2-morpholinoethyl)carbodiimide metho-p-toluenesulfonate, Sigma-Aldrich) reagent based on protocol published by Carlile et al (34). Specifically, rRNA pellet has been dissolved in 20 uL volume of 0.4 M CMCT in BEU buffer (50 mM Bicine (pH 8.5), 4 mM EDTA (pH 8.0), 7 M Urea), and incubated at 37 °C for 1h. Then, rRNA has been purified from excess of CMCT using 0.5 mL 30 kDa Amicon Ultra filter with two rounds of exchange to RNase-free water. The recovered 30-40 uL of rRNA were combined with an equal volume of 100 mM sodium carbonate-bicarbonate buffer (pH=10.5) and incubated at 50 °C for 2h. Following additional two rounds of 30 kDa Amicon buffer exchange to RNase-free water, CMC-derivatized rRNA has been freeze-dried and subjected to the RNase digestion. CMCT protocol used resulted in 60-90% efficient labeling of all rRNA pseudouridines, without significant number of off-target identifications. A single CMC chemical tag adds 251.2 Da to the mass of underivatized nucleolytic RNA, and furthermore increases its retention of the reverse-phase column, thus permitting specific detection of pseudouridines in addition to rRNA methylation.

LC-MS data were acquired on Agilent Q-TOF 6520 ESI instrument coupled to the Agilent 1200 LC system. rRNA digestion fragments were chromatographically resolved on XBridge C18 column (3.5 µM, 1×150 mm, Waters) using 15 mM ammonium acetate (pH = 8.8) as mobile phase A and 15 mM ammonium acetate (pH = 8.8) in 50% acetonitrile as mobile phase B. 50 min long linear gradient was constructed: 1-15% of B over 40 min (corresponds to elution of underivatized RNA), followed by 15-35% of B over 10 min (corresponds to elution of CMC-derivatized RNA). Negative ionization data were recorded over the 400–1700 m/z range. Pairs of the co-eluting ^14^N- and ^15^N-labeled fragments were identified using their predicted m/z values. MS peak profiles were extracted, averaged over a 0.1 min wide window, and quantitatively fit to their theoretical isotope distribution using Isodist (35). The ratios of ^14^N- and ^15^N-peak amplitudes were used to calculate modification fractions for individual rRNA modifications, after they were normalized to the ratios for unmodified 16S or 23S rRNA transcripts (33).

## RESULTS and DISCUSSION

### Assembly of the Ribosome in iSAT is near-native

In the iSAT reaction, the functional 70S ribosome is continuously synthesized, and progress of the assembly is monitored by translation of a GFP reporter (Figure 1A). The fluorescence signal starts to rise after a lag of about 20 minutes. As shown in Figure 1B, when the purified 70S ribosomes were added to the reaction instead of the plasmid encoding rRNA and purified total ribosomal proteins, the fluorescence signal rises without a lag, indicating that the signal lag observed in the iSAT reaction can be attribute to the assembly of the ribosome. The fluorescence signal reaches a plateau after 2 hours (Figure 1B), which is likely due to exhaustion of the ATP supply (36). In comparison with the ribosome profile for the 70S ribosome control, the sucrose gradient profile of 4-hour iSAT reaction (Figure 1C) indicates the presence of a pre-50S peak in addition to the 30S peak and a much lower 70S peak, which can be attributed to the limitations of the *in vitro* environment, such as exhaustion of materials or unbalanced production of rRNA with respect to r-proteins. The pre-50S peak can be characterized to provide information about the ribosome assembly process.

**Figure 1.**
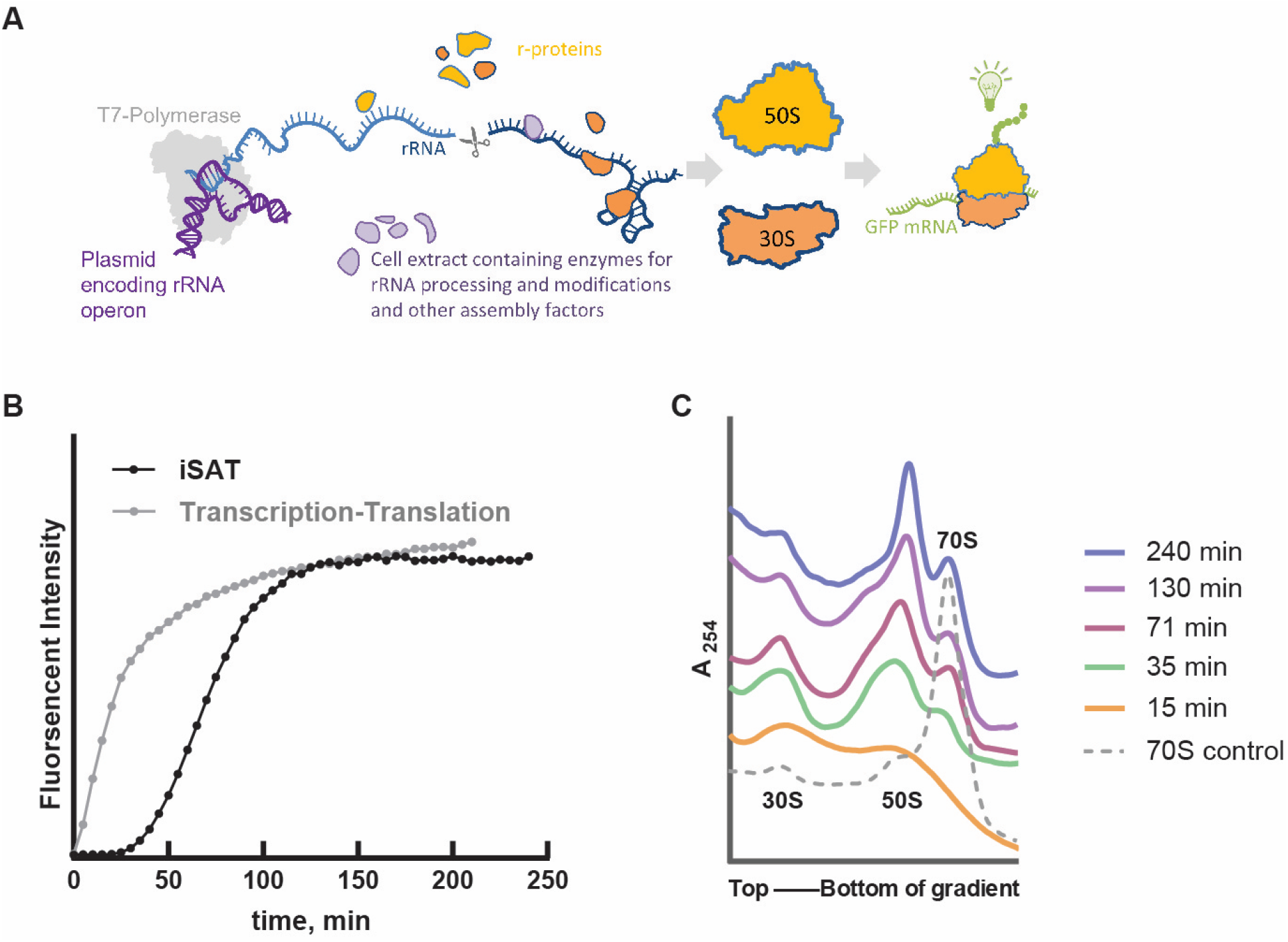
Time course of integrated rRNA synthesis, ribosome assembly, and translation (iSAT). **(A) Schematic of iSAT reaction**. In an iSAT reaction, plasmids encoding rRNA and a reporter mRNA are transcribed by T7 polymerase. The rRNA is processed and modified by enzymes in an *E*. coli cell extract, and ribosomal proteins derived from purified ribosomes bind to complete the ribosomal subunits. Newly assembled ribosomes engage in the translation of the reporter protein (GFP) to produce a readout of production of functional ribosomes. **(B) Fluorescence readout of iSAT**. The time course of GFP fluorescence is shown for a standard iSAT reaction (black), compared to a control reaction where intact ribosomes were directly added into the reaction (gray). The delay of fluorescence signal in the iSAT reaction is due to assembly of sufficient ribosomes to produce GFP. **(C) Ribosome profile of iSAT time course reactions. Five parallel** iSAT reactions were quenched at sequential time points, and the ribosome profiles of the iSAT were analyzed using sucrose density gradient ultracentrifugation. In general, both the size and abundance of ribosome precursors increases over time.

The completed 70S ribosomes produced during iSAT were analyzed for their protein composition and rRNA modifications using quantitative mass spectrometry (Figure S1). Most ribosomal proteins were present in stoichiometric amounts compared to purified 70S ribosomes from MRE600, except that bL33 was reduced by 53%. Post-transcriptional modifications (PTMs) of ribosomal RNA play a critical role in the assembly process, with consequences for RNA folding, ribosome activity and susceptibility to metabolites (37-39). Remarkably, we found that most nucleotide modification sites on both 16S and 23S rRNAs were properly installed in the iSAT reaction (Table S1), which supports the idea that the recruitment of rRNA modification enzymes from the cell extract during the iSAT reaction resembles the native process *in vivo*.

### Time-resolved *in vitro* assembly of the LSU

The iSAT reaction is both continuous and asynchronous, and it is likely that there is a pool of assembling ribosomes over the time course of the reaction, with accumulation of completed ribosomes over time. To capture and examine an array of the ribosome intermediates of various stages, the iSAT reaction was quenched at a series of 15, 35, 71, 130, and 240 minute timepoints, from the very beginning stage where no fluorescence signal was detected, to the very late plateau stage (Figure 1B). The sucrose gradient profiles of the reaction mixture showed that the particle size generally increased with time (Figure 1C), implying the presence of more immature pre50S particles at earlier time points, and the opportunity to examine a range of intermediate structures.

### Iterative ab initio subclassification reveals new LSU assembly intermediates

In order to analyze the highly heterogenous cryo-EM dataset from our iSAT reaction, we developed an iterative sub-classification workflow based on cryo-SPARC (Figure 2). To reconstruct as many classes as possible, especially the minor classes, the particles extracted from the motion-corrected micrographs of all five time-course datasets were pooled together for the processing. Two rounds of 2D classification were performed to remove the particles that are obviously not pre-50S intermediates, such as 30S and 70S particles. Then the remaining particles were iteratively sub-classified using *ab initio* reconstruction, until there are not enough particles in a given subclass to provide a 10 Å reconstruction. The final maps resulting from the iterative sub-classification were refined by homogeneous refinement. Using this workflow, a set of thirteen distinct maps was identified from all the time course datasets, as shown in Figure 2A. The resolutions of the maps vary from 4.53 Å to 8.83 Å, primarily depending on the number of particles in each class.

**Figure 2.**
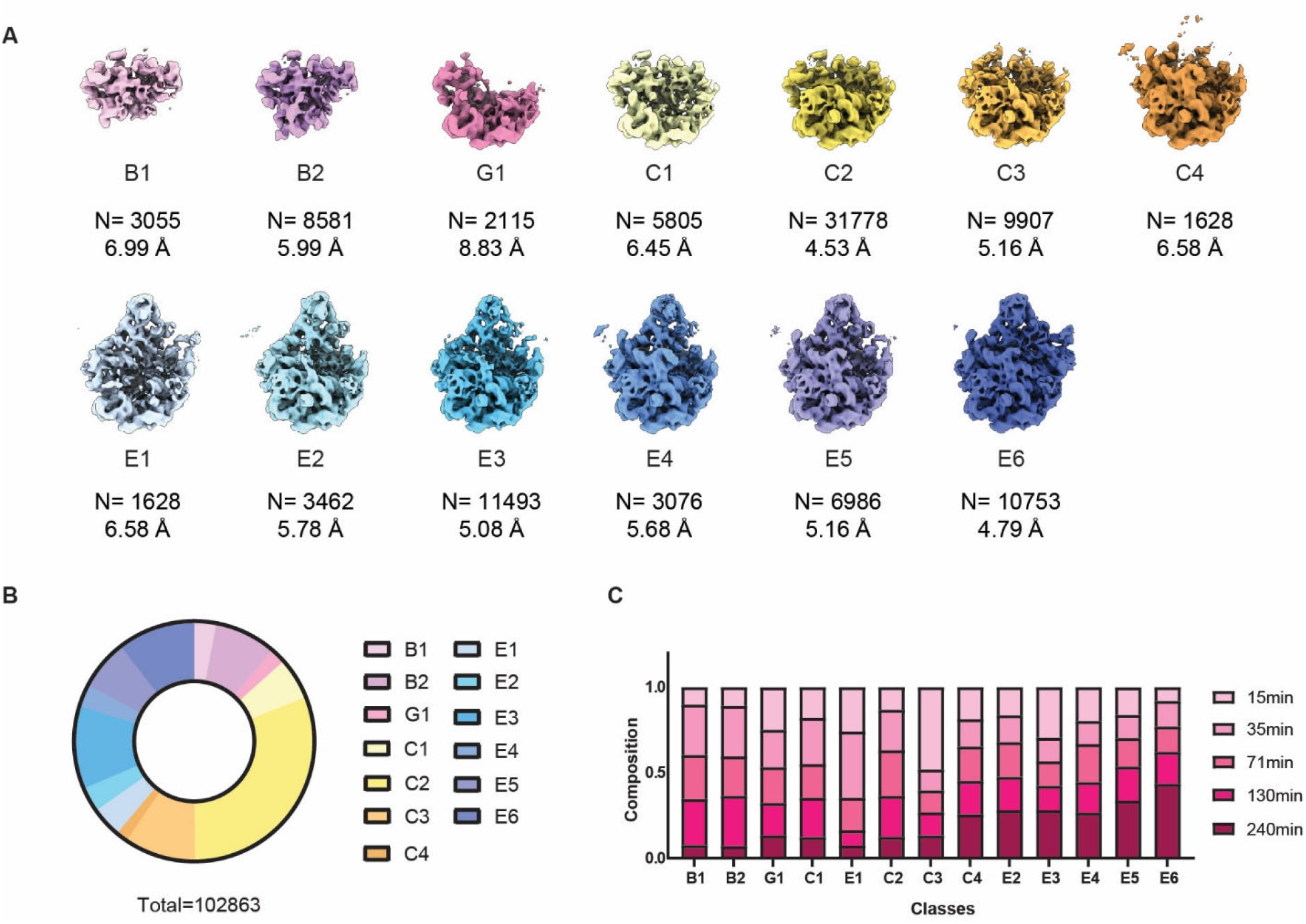
Distribution of 50S intermediates in the iSAT reaction time course. **(A) Density maps for 50S intermediates**. Thirteen 50S **intermediates** were obtained by heterogeneous subclassification from the iSAT reaction time course, where all timepoints were combined prior to analysis. The thirteen 50S precursors are named by the major classes they belong to, and are generally ordered from immature to mature. The number of particles contributing to each class is given along with the final resolution of the map. **(B) Particle class distribution among the 50S intermediates**. The particle distribution among 50S precursors is calculated according to the number of particles reconstructing each class (Table S3). **(C) Temporal composition of 50S intermediates**. The contribution of each timepoint for each 50S **intermediates** class was calculated and plotted in the bar plot. The number of the particles was firstly normalized in each dataset of the five timepoints. And the ratio of the particles from different timepoints was calculated in each class. In general, the less mature classes have a higher contribution from the earlier timepoints, and the more mature classes have a higher contribution from the later timepoints. The production of ribosomes is continuous during the iSAT reaction, and it is expected that contributions from all classes could be observed at any timepoint.

The thirteen iSAT intermediates were compared with the previously identified B, C, D, and E-class intermediates of the bL17-depletion dataset (13) (Figure S3A), revealing four C classes, no D class, and six E classes in the iSAT intermediate ensemble. Notably, two very early B classes and a new class, named G, were identified. The fraction of total particle numbers for each class was determined, and about one half (48.1%) of the particles belong to C class, while the E class occupies 39.1%, while the B and G classes account for 10.8% and 2.1%, respectively, as shown in Figure 2B. The contribution of each timepoint to each of the classes (Figure 2C) indicates a trend, where the more mature classes originate from the particles of later timepoints. To summarize, the iSAT intermediate library covers multiple classes from early to late stages, which can provide comprehensive information about the entire course of 50S assembly.

In 2018, Nikolay et. al reported five structures (State 1 to State 5) from *in vitro* reconstituted 50S precursors using the Nierhaus reconstitution protocol (17). Figure S3B compares those 5 precursors with the 13 iSAT intermediates in the present work by hierarchical analysis of molecular weight differences. According to the clustering, B1, B2, C1 and G1 from the iSAT dataset were clearly separated from the more mature classes, indicating that iSAT intermediates cover the earlier assembly stage. States 1 and 2 were closest to the C class. Examination of the maps revealed that State 1 appears to be a stage between C1 and C2, while State 2 is very similar to C2, which is the most abundant class among the iSAT intermediates. Considering that State 2 was part of the 41S particles trapped in the first step of the classical *in vitro* reconstitution (44 °C, 4 mM Mg^2+^), C2 class and State 2 could represent a thermodynamically stable stage during the ribosome assembly, where many intermediate particles were trapped. State 3 is similar to E2, while State 3 resolved a more completely formed L1 stalk. State 4 and State 5 are close to E5, which represent the late assembly stage. Overall, the iSAT intermediates span similar states to the *in* vitro reconstitution intermediates, provide more information of early stages, and increase the diversity of late structures.

### Segmentation of the set of Maps Identifies Cooperative Assembly Blocks

To quantitatively analyze the structural features of each intermediate maps, the mature 50S structure was divided into 139 segments based on the RNA secondary structures and ribosome proteins (13, 31) (rows in Figure 3A). Taking these segments as the reference, the occupancy values for each segment can be calculated for the set of intermediate maps (columns in Figure 3A). The profile of each segment was analyzed among all the classes by quadrant analysis, as described in Methods. The segments that have the same behaviors are sorted out as a cooperative assembly block with correlated values of the occupancy matrix across the classes. From this, 14 Blocks were defined and named based their most significant structural or compositional features (Figure 3B and Table S2).

**Figure 3.**
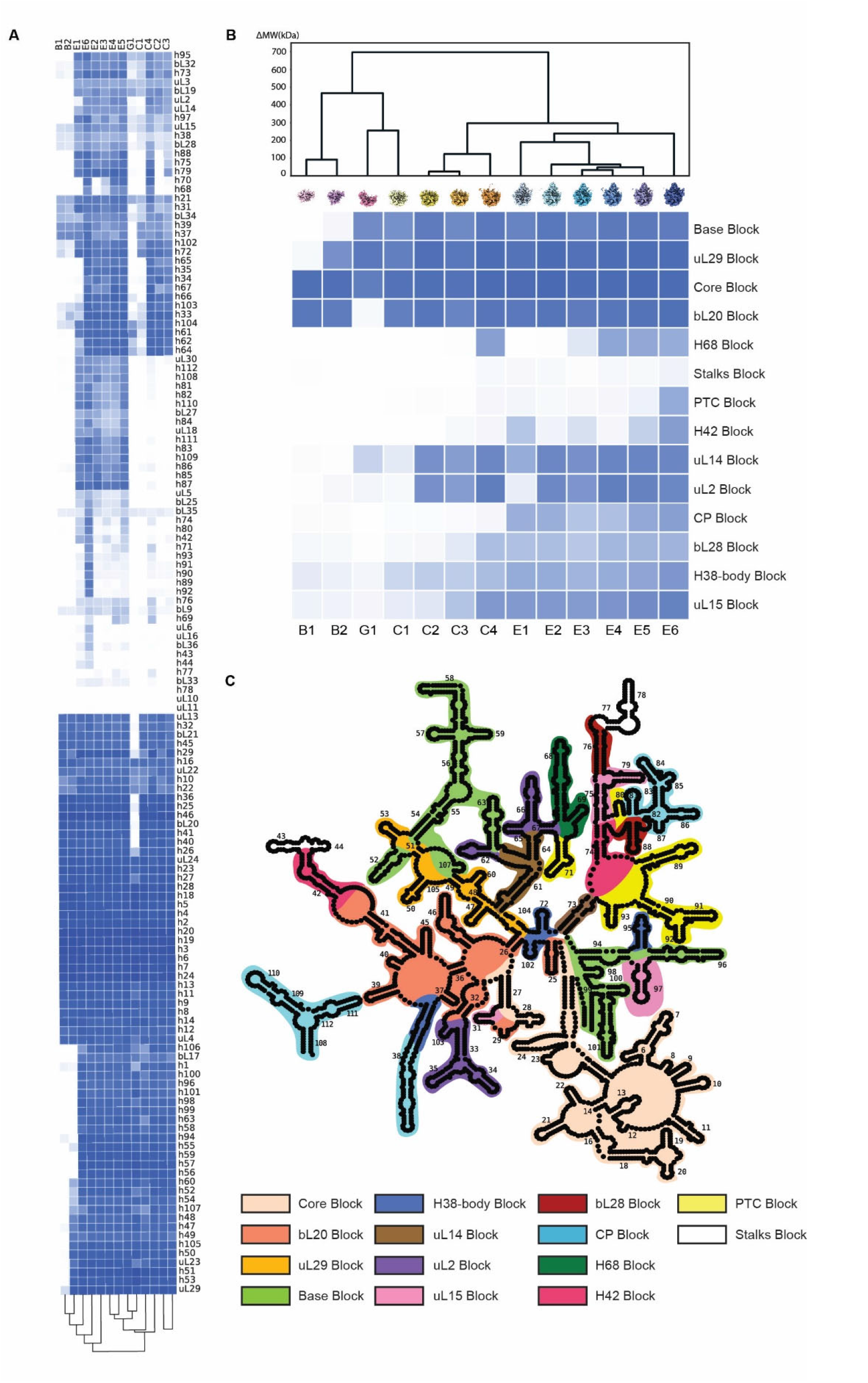
Occupancy analysis of helix, protein, and assembly blocks. **(A) Electron density occupancy of rRNA helices and ribosomal proteins**. The occupancy values are calculated as described in Methods and are used to analyze the correlation of the assembly elements as rRNA helices and ribosomal proteins for each of the 13 50S intermediates density maps. The elements correlated to each other are determined using hierarchical clustering, and are defined as an assembly block in part (B). **(B) Electron density occupancy of the assembly blocks**. The occupancy values are calculated as described in Methods and are used to analyze the dependencies between the assembly blocks. The blocks are named for prominent structural features, protein association, or RNA helical elements. The dendrogram of hierarchical clusters were calculated according to the Euclidean distance matrix (in molecular weight, kDa) among the density maps of 50S **intermediates. (C) rRNA composition of the assembly blocks. The** rRNA helices in the 23S rRNA secondary structure map are colored according to the assembly blocks.

The rRNA composition of each assembly block is shown in Figure 3C. The rRNA elements in most of the blocks are either contiguous in sequence and secondary structure, or are involved in tertiary interactions. The occupancy of the elements within blocks is correlated across the datasets, indicating they can assemble as modules, or cooperative folding blocks. A few exceptions include H38-body Block, the uL15 Block and the H42 Block. In these 3 blocks, rRNA fragments are temporally, but not spatially related, meaning that they assemble at the same stage, but may not have direct interactions. Notably, the long H38 element can be divided into two parts, one of which belongs to the CP Block and the other to theH38-body Block, respectively. This finding indicates that H38 may function as a “bridge” to connect the central protuberance to the main body of the 50S subunit (Figure S7).

### The Assembly Blocks Fold Following Certain Dependencies

By analyzing the order of appearance of blocks, we determined the dependencies between the blocks (Figure 4 and Figure S5). Briefly, if Block B is never observed without Block A, we conclude that Block B has a dependency on Block A, and that the assembly of Block B happens after the assembly of Block A, while Block A is capable of folding without Block B. By following the dependencies among blocks, a putative set of 22 possible intermediate structures can be identified. However, not all the putative structures are stable and abundant enough to be resolved in our iSAT intermediate library. The observed 13 structures are a subset of the 22 possible structures based on the observed dependencies, and likely represent thermodynamic waypoints during assembly. The structures that are consistent with the block dependencies but are not observed may never form, or be transient species that do not accumulate during the iSAT reaction.

**Figure 4.**
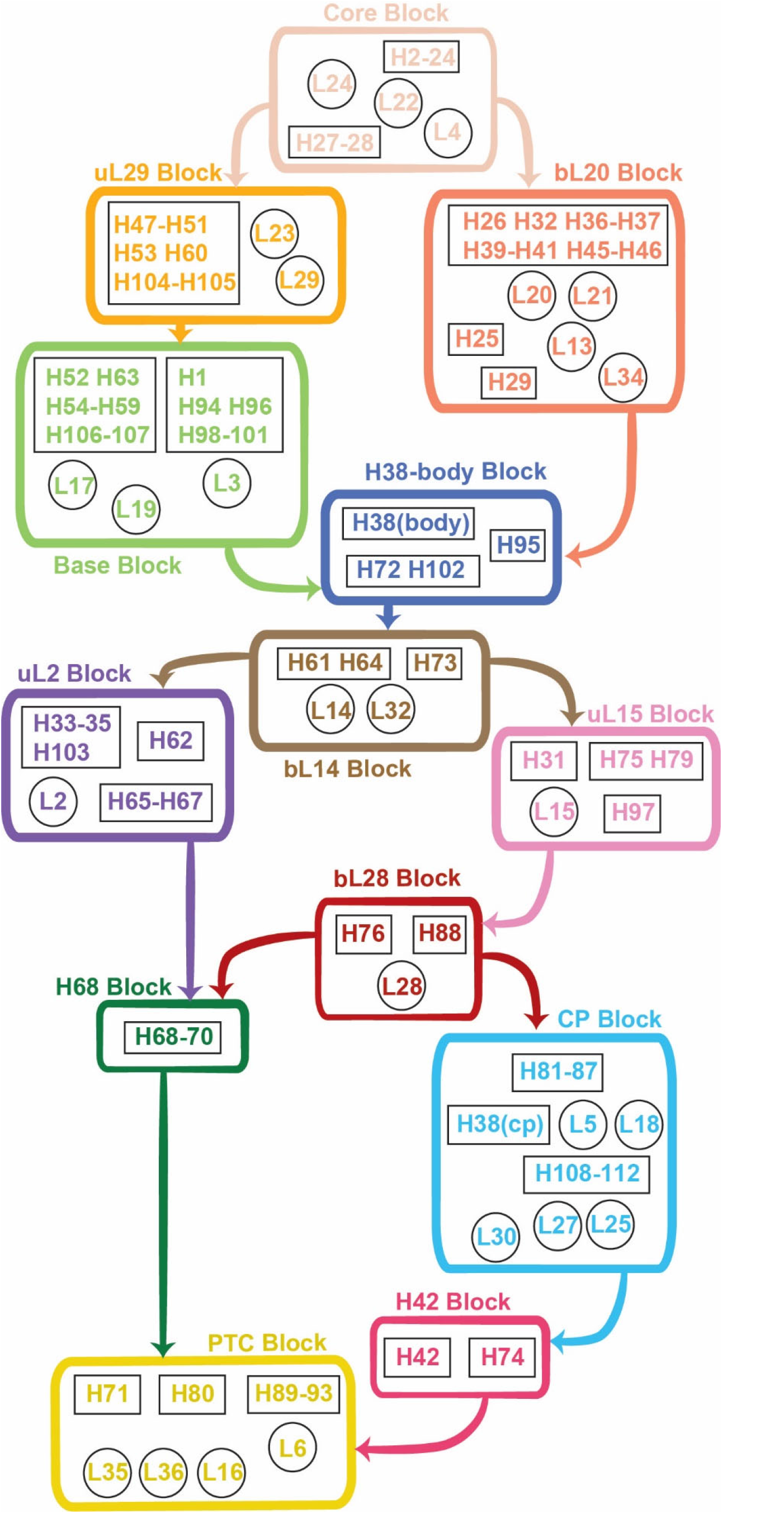
Complete 50S assembly map including RNA and protein elements. The assembly blocks and their downstream dependencies are colored according to Figure 3C. Block dependencies were determined using a quadrant analysis as described in Methods and are shown as bold colored arrows. The rRNA helices are outlined in black rectangle boxes, while the ribosomal proteins are in the black circles.

The dependency map shown in Figure 4 illustrates not only the dependencies between the assembly blocks, but also some independencies among the blocks. For example, after the formation of the Core Block, the uL29 Block and bL20 Block can assemble onto the Core Block independently. With these dependencies and independencies among the assembly blocks identified, we learned that, firstly, some of the rRNA helices and the ribosomal proteins that interact strongly with each other and assemble as modules, defined as cooperative assembly blocks in this work. Second, the assembly of some cooperative blocks depend on the assembly of other blocks in a defined way that may limit the complexity of the assembly pathway. Thirdly, in addition to the dependencies, the assembly of some blocks are independent, providing parallel pathways ensuring the high efficiency of the assembly.

### Updated Nierhaus assembly map including the assembly block hierarchy

The ribosomal protein composition of each block is labeled on the Nierhaus map, as shown in Figure 5. The thermodynamic binding dependencies from the original Nierhaus map are retained (19), but the protein positions in the map were re-organized. The horizontal position was arranged according to contacts to the 23S rRNA helices. Some r-proteins have contacts with the rRNA helices in more than one assembly block, but primary binding can be assigned to the rRNA helices where the r-protein was first resolved in the series of intermediates. The vertical position was determined by the association of the r-proteins with the assembly blocks, according to the dependencies among the assembly blocks. This update confirms the validity of the original thermodynamic dependencies, and clearly conveys the 5’-3’ progression of the assembly process via cooperative assembly blocks. As shown in Figure 5, the assembly starts from the left top, with the initiating proteins interacting with the 5’ end of the 23S rRNA. As the assembly proceeds, the binding of the proteins with the 23S rRNA moves gradually to the 3’ end. In brief, the updated Nierhaus assembly map illustrates more detailed information about the interactions between the r-proteins and the rRNA helices, which also provides evidence for the co-transcriptional direction for assembly.

**Figure 5.**
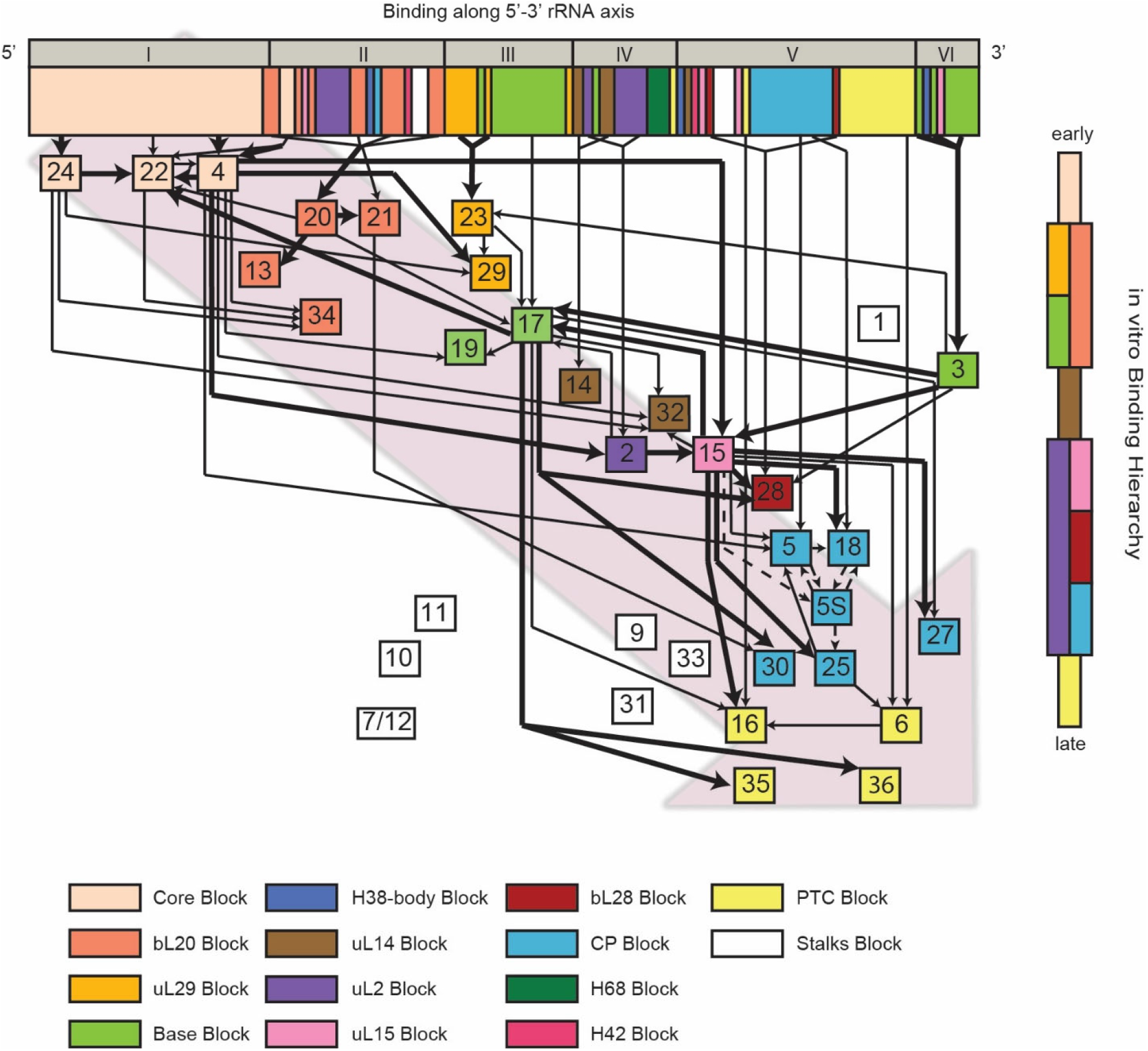
Updated Nierhaus assembly map including assembly block hierarchy. The original protein dependencies from the Nierhaus assembly map are shown as thick and thin black arrows, for strong and weak dependencies. The 23S rRNA bar is annotated both with the domain designation and colored according to the iSAT assembly block. The order of the block dependencies in Figure 4 is shown at the right to indicate early to late progress. The positions of the ribosomal proteins are rearranged horizontally according to the position of the interaction rRNA helices in the same block, and rearranged vertically according to the dependencies of the assembly. Remarkably, the basic structure of the original Nierhaus assembly map is consistent with the 5’-to 3’ co-transcriptional direction of assembly and with the newly observed block dependencies.

Several proteins located at the high flexible stalks were not well-resolved in our intermediate library, such as uL10, uL11, and bL7/12. Only a small part of bL9 interacting with H76 was resolved, suggesting the point of anchoring of the dynamic L9. In addition, bL33 was not observed with significant occupancy in the resolved thirteen intermediates, consistent with the mass spectrometry result (Figure S1A). As Davis et al. reported, bL33 has dependency on bL17, but no reduction of bL17 was detected in our intermediate library, so the absence of bL33 can be possibly due to an unknown dependency on other ribosomal proteins or assembly factors.

### Early pre50S Intermediates Reveal an Assembly Core

The Core Block, as shown in Figure 6A, is the intersection of the density for the B1 and G1 classes. The assembly core reported in this work is the earliest pre50S structure reported so far. The structure of a similar assembly core was also identified in another dataset obtained from the deletion strain DdeaD (submitted), which confirms the important role of this assembly core for the initiation of the assembly of LSU. The Core Block is composed of Helices 2-24, Helices 27-28, and three ribosomal proteins: uL4, uL22, and uL24 (Figure 6B). All these helices are at the 5’ end of the rRNA transcript, consistent with the hypothesis of co-transcriptional ribosome assembly (18, 40, 41).

**Figure 6.**
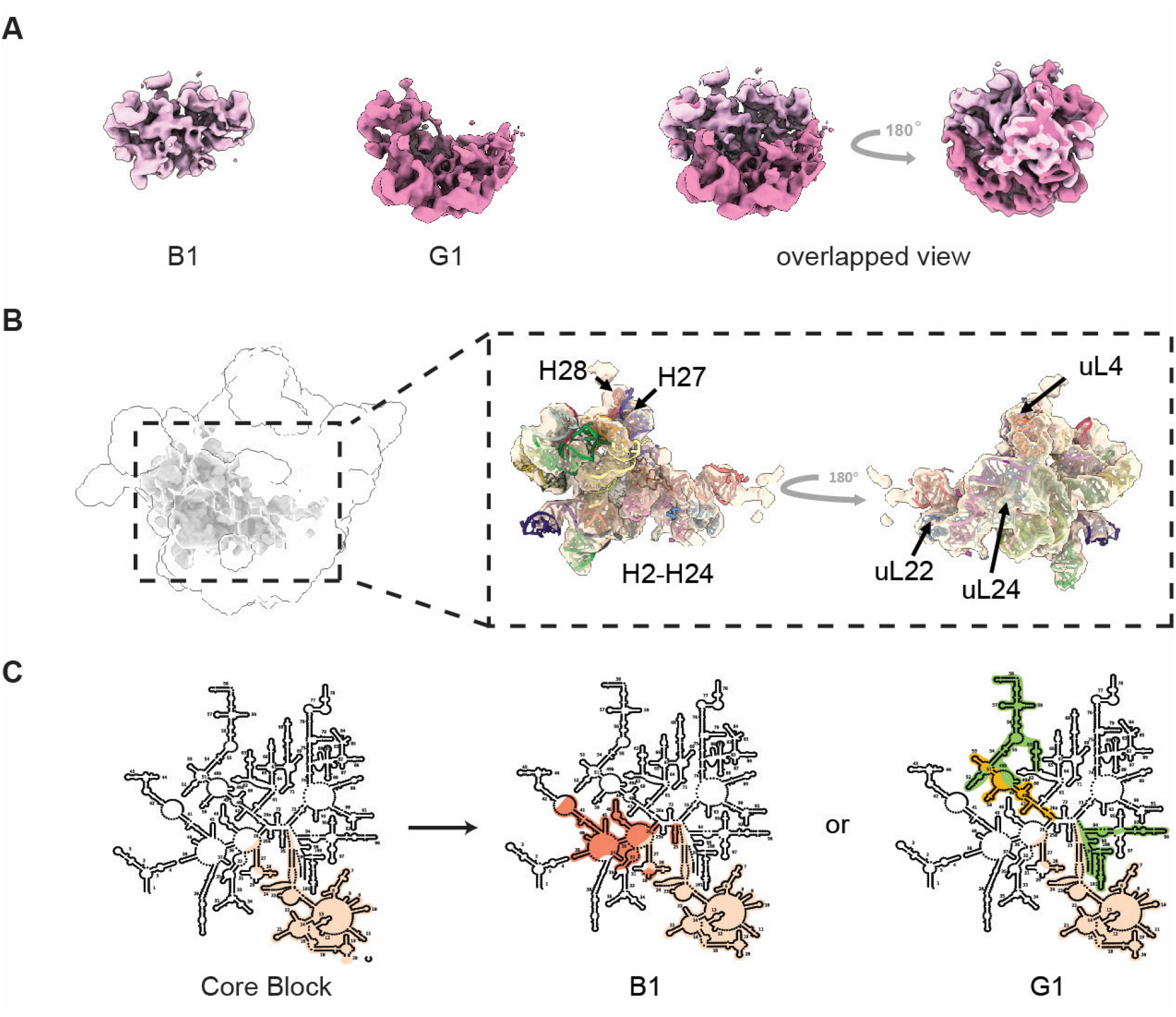
Evidence for parallel steps very early in assembly. **(A) The overlapped view of 50S intermediates B1 (upper) and G1 (lower)**. The density region in common between the B1 and G1 classes corresponds to the Core Block. **(B) Parallel pathway early in assembly**. Formation of the B1 class and the G1 class occurs by consolidation of either domain II (for B1) or domains III/VI (for G1). The rRNA secondary structure maps are colored according to the blocks, indicating two parallel pathways after the folding of Core Block. The colors of rRNA helices are correlated to the previous figures. **(C) Core Block**. The position and structure of the assembly Core Block are displayed. The rRNA helices and ribosomal proteins UL4, uL22, and UL24 in Core Block are labeled, consisting with formation of domain I as the earliest step in assembly (Figure 5).

The structural difference between B1 and G1 class implies two independent assembly pathways after the formation of the Core Block (Figure 6C). On the pathway towards B1, the bL20 Block assembles on the Core Block, which is missing in G1. The bL20 Block includes other early ribosomal proteins such as bL20, bL21, bL34 and uL13, and the rRNA fragments in bL20 Block are subsequently transcribed after the helices in Core Block. On the other pathway, G1 completes the docking of Base Block and uL29 Block. To summarize, the existence of Core Block provides a key insight into the initiation of the assembly of the ribosome at the 5’-end.

### Large Ribosomal Subunit Assemble by Parallel and Modular Pathways In Vitro at Both Early and Late Stages

The analysis of block composition for each class suggests parallel and modular assembly pathways (Figure 7 and Figure S6). The B and G classes reveal two parallel early assembly pathways. After the early assembled Core Block binding with bL20 Block to form B1, uL29 Block can assemble to form B2. The forming of bL20 Block and lack of Base Block is the feature of B class in iSAT intermediates library, while G class is just the opposite. Through either B or G pathway can reach C1 class, which contains all the Blocks in B and G classes.

**Figure 7.**
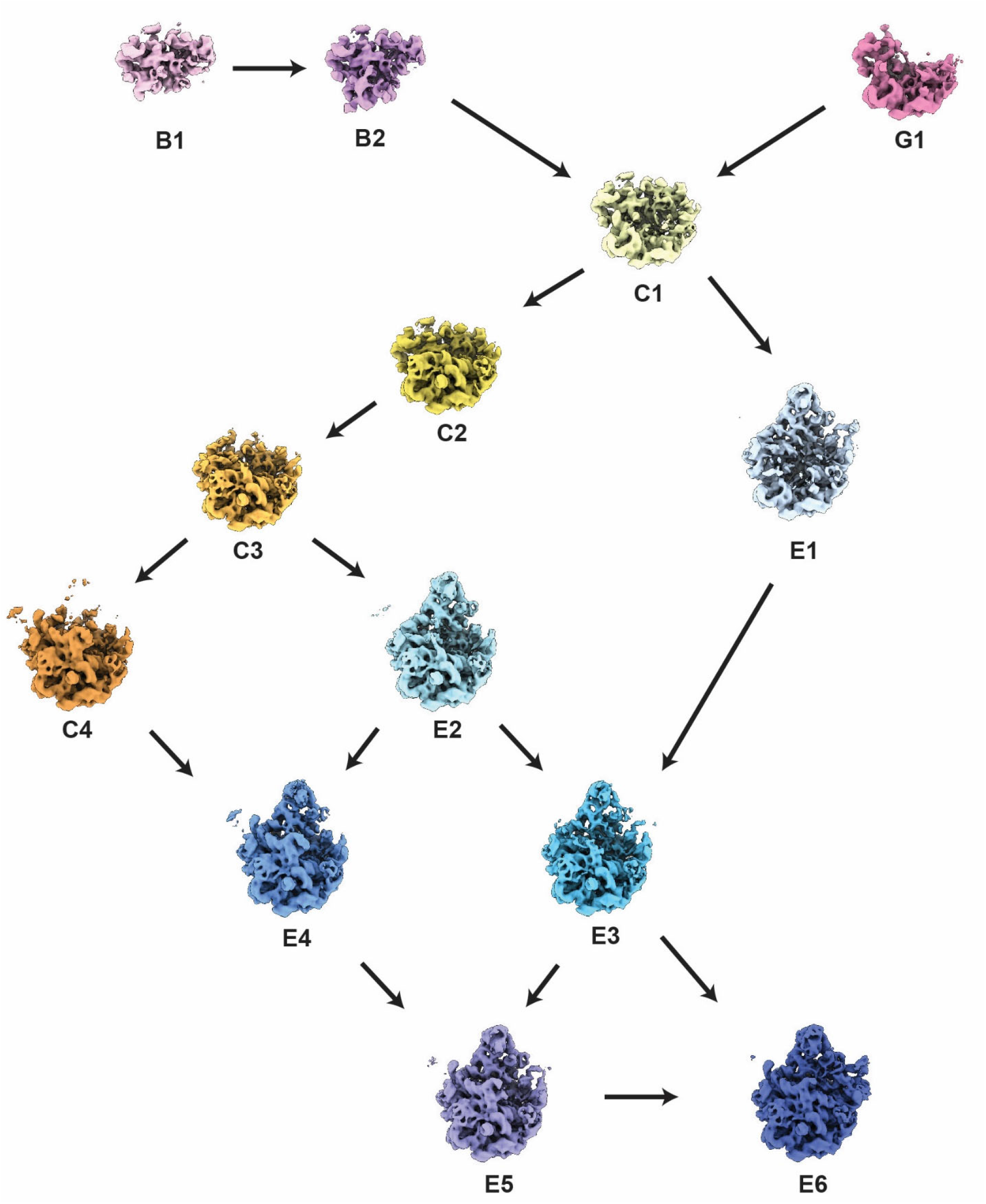
Organization of 50S intermediates into an assembly pathway. The thirteen reconstructed 50S **intermediates** from iSAT reaction time course are organized according to the dependencies of the assembly blocks. The arrows indicate the minimal folding steps required to connect the set of **intermediates**.

The H38-body Block also forms in the C1 class. From the C1 class through the C4 class, more and more H38 density is resolved (Figure S7), indicating the forming progression of this long bridge helix. The most significant feature of the C2 class is the formation of Blocks at the “belly” of the subunit: uL14 Block and uL2 Block. After that, the C3 class forms the uL15 Block, which is another key block required for docking of the CP Block. According to the Nierhaus map and Figure 5, uL15 has multiple interactions with the proteins in CP Block (19), and is a critical protein for the initiation of the assembly of the CP (42). With the uL15 Block assembled, the C3 class is ready for docking of the CP Block.

The assembly at late stage is more parallel. From the C3 class, either H68 Block or CP Block is ready to assemble. In the C3-C4-E4 pathway, the H68 Block first assembles to form the C4 class, then the CP Block assembles to form the E4 class, while the C3-E2-E4 pathway is an alternative sequence of transitions. The increased density of the H42 Block in the E3 class is the primary distinction from the E4 class. With all of the above blocks assembled, the E5 class can support formation of the PTC block to reach the E6 class, which is the most mature class resolved in the iSAT intermediate library.

The E1 class exhibits the lack of the uL2 and H68 Blocks, indicating another parallel pathway differing from the late-stage pathways mentioned above. The closet precursor of the E1 class is the C1 class, which may be due to some transitional intermediates that are not resolved in our iSAT intermediates library. The presence of E1 indicates the high parallel and modular feature of 50S assembly at late stage.

In summary, the iSAT reaction provided a wealth of information about the intermediates that accumulate during assembly of the 50S subunit under near-physiological conditions. The intermediates share common features with previous datasets resulting from perturbed growth in cells or *in vitro* reconstitution (7, 13, 17). In addition, a significant series of novel early intermediates were identified that provide the earliest known intermediates to date for 50S assembly, The features of parallel and sequential assembly steps appear to be ubiquitous on the ribosome assembly landscape, although the assembly is generally consistent with the expected co-transcriptional direction of rRNA synthesis. The flexibility of the iSAT reaction conditions will provide a powerful tool to further probe the mechanistic roles of individual assembly factors that are critical for efficient assembly under a variety of conditions in cells.

## Acknowledgements

This work was supported by grants from the National Institutes of Health GM136412 and GM053757 to JRW.

## Notes

### Competing Interest Statement

The authors have declared no competing interest.

