## Supplementary Material for "Near-Physiological in vitro Assembly of 50S Ribosomes Involves Parallel Pathways"

**This supplement contains:**
Supplementary Figure S1 to S6
Supplementary Table S1 to S3

### SUPPLEMENTARY FIGURES

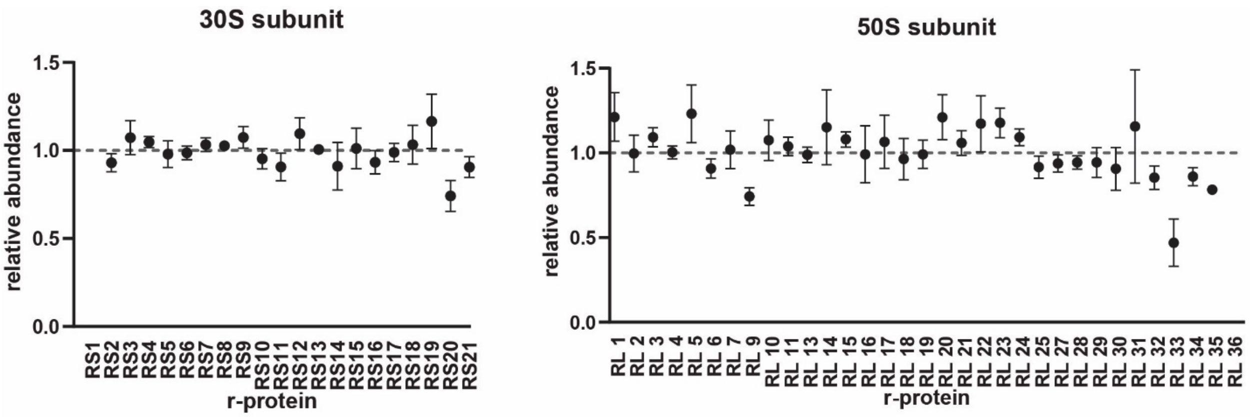


**Figure S1. Mass spectrometry analysis of 70S ribosomes from iSAT.**

r-protein composition in 30S and 50S ribosomes assembled from using the iSAT reaction quantified by mass spectrometry compared to $^{15}N$ 70S ribosomes, as described in Methods.

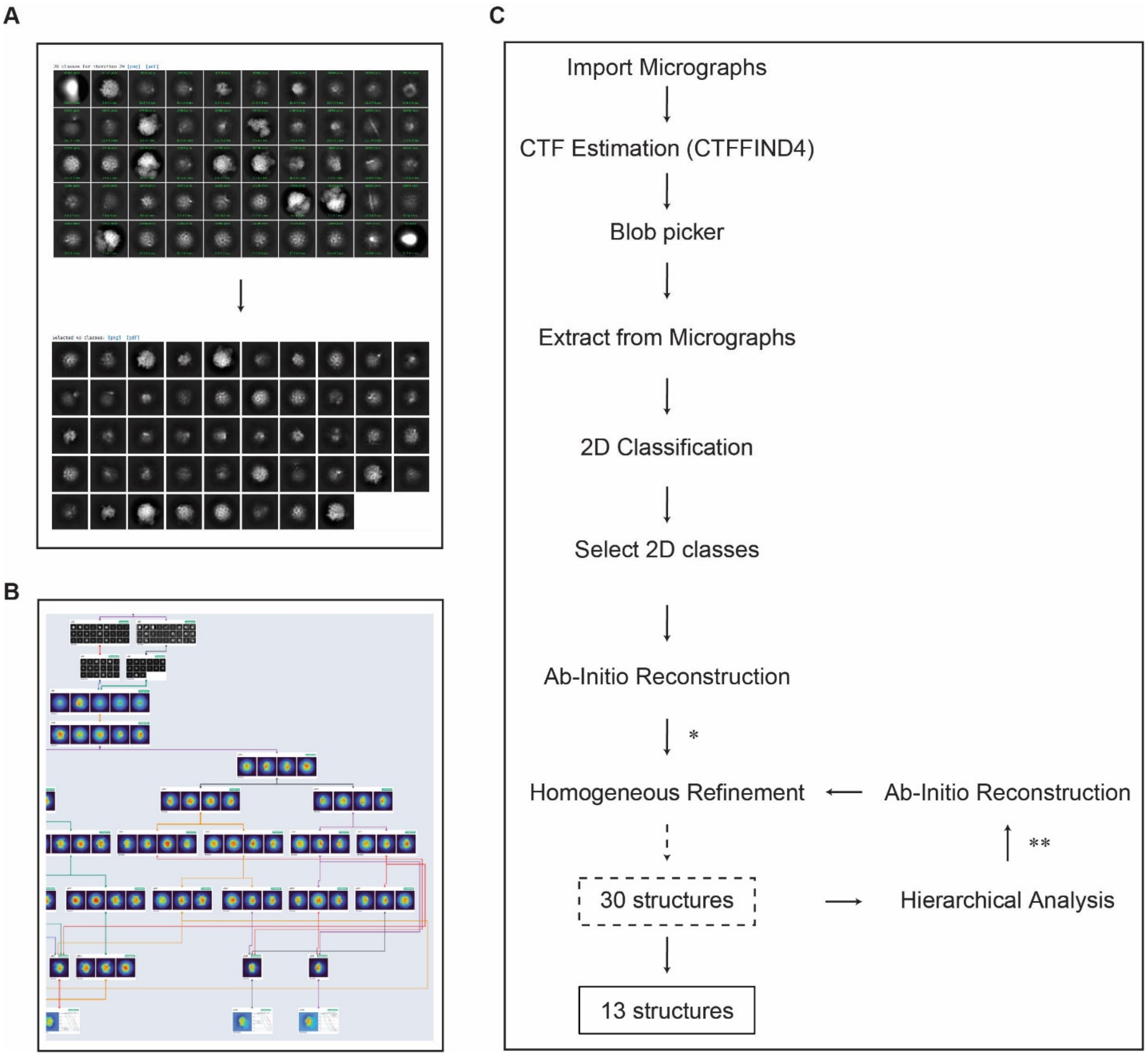


**Figure S2. Workflow for iterative subclassification of heterogeneous particles from cryo-EM data.**

**(A)** The screenshot of 2D classification and 2D class selection performed in cryoSPARC. **(B)** The screenshot of iterative subclassification by ab-initio reconstruction in cryoSPARC. **(C)** The iSAT time course dataset was analyzed according to the workflow indicated here. More details are described in Method. *: The particles are iteratively subclassified by ab-initio reconstruction. When the resolution of reconstructed class is less than 10 angstroms, the subclassification of this class will be finish, and homogeneous refinement of this class will be performed. **: According to the results of hierarchical analysis, the similar classes will be combined through consensus ab-initio reconstruction and homogeneous refinement. The combined classes will be checked by hierarchical analysis to determine if more combinations are needed.

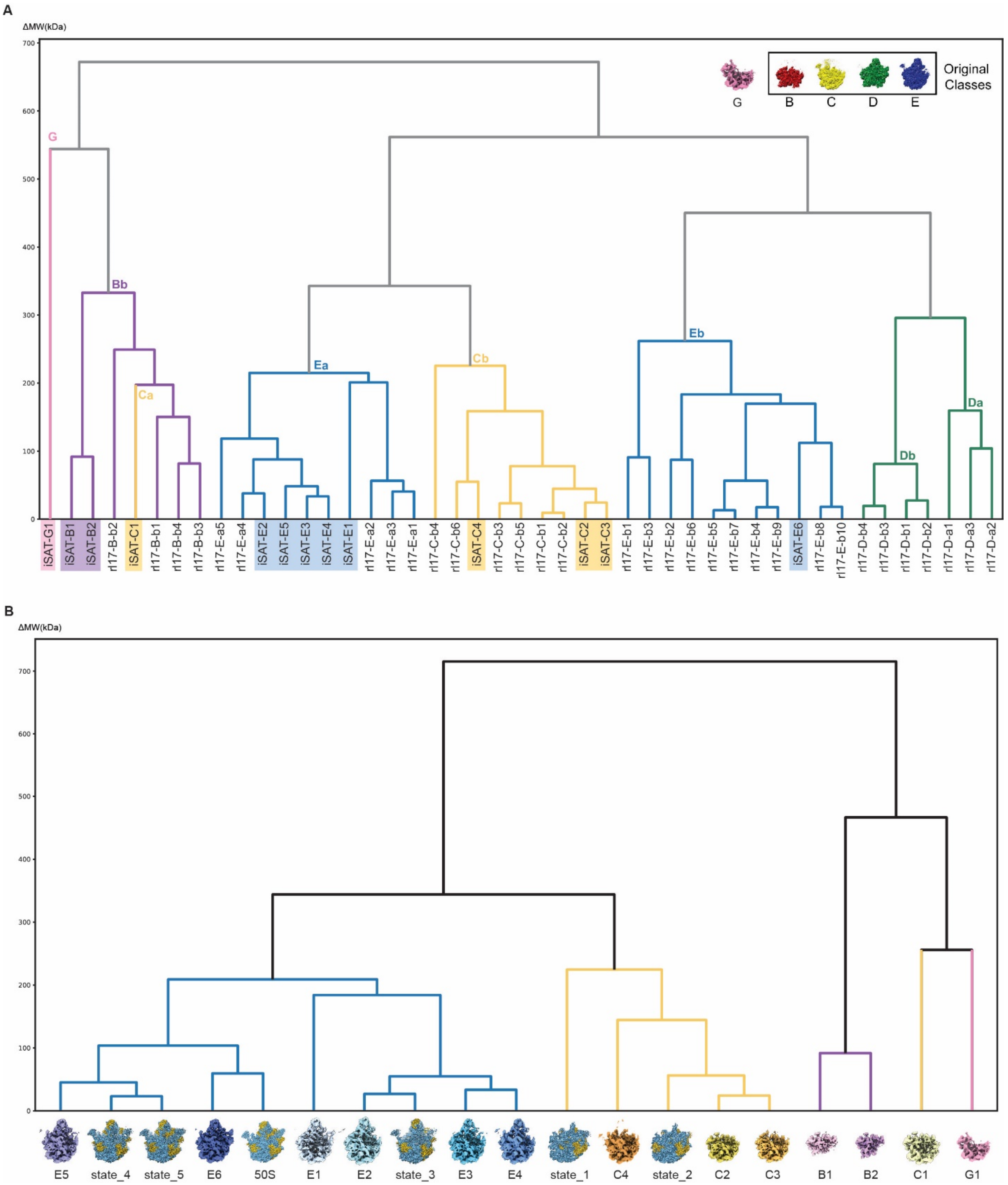
A
ΔMW(kDa)
700
600
500
400
300
200
100
0
Original Classes
G
B
C
D
E
G
Bb
Ca
Ea
Cb
Eb
Da
Db
iSAT-G1
iSAT-B1
iSAT-B2
rl17-B-b2
iSAT-C1
rl17-B-b1
rl17-B-b4
rl17-B-b3
rl17-E-a5
rl17-E-a4
iSAT-E2
iSAT-E5
iSAT-E3
iSAT-E4
iSAT-E1
rl17-E-a2
rl17-E-a3
rl17-E-a1
rl17-C-b4
rl17-C-b6
iSAT-C4
rl17-C-b3
rl17-C-b5
rl17-C-b1
rl17-C-b2
iSAT-C2
iSAT-C3
rl17-E-b1
rl17-E-b3
rl17-E-b2
rl17-E-b6
rl17-E-b5
rl17-E-b7
rl17-E-b4
rl17-E-b9
iSAT-E6
rl17-E-b8
rl17-E-b10
rl17-D-b4
rl17-D-b3
rl17-D-b1
rl17-D-b2
rl17-D-a1
rl17-D-a3
rl17-D-a2
B
ΔMW(kDa)
700
600
500
400
300
200
100
0
E5
state_4
state_5
E6
50S
E1
E2
state_3
E3
E4
state_1
C4
state_2
C2
C3
B1
B2
C1
G1

**Figure S3. Comparison of iSAT particle maps with other structural datasets using hierarchical clustering.**

**(A) Clustering organizes the set of iSAT 50S precursors and the set of bL17-independent 50S precursors into similar groups.** The Euclidean distance matrix (in molecular weight, kDa) was calculated among density maps, and the dendrogram resulting from hierarchical clustering is displayed, with 5 main class branches colored accordingly. The color-highlighted labels represent the density maps from iSAT time course dataset, while others are from bL17-independent dataset. C1 of iSAT time course dataset is classified into C class according to the hierarchical analysis result with a larger dataset (data not shown) **(B) Hierarchical clusters of iSAT 50S precursors and 50S precursors of Nikolay et al.** The dendrogram is drawn in the same principle described in (**A**). The maps labeled as state_1 to 5 and 50S_rec are from the dataset of Nikolay et al.

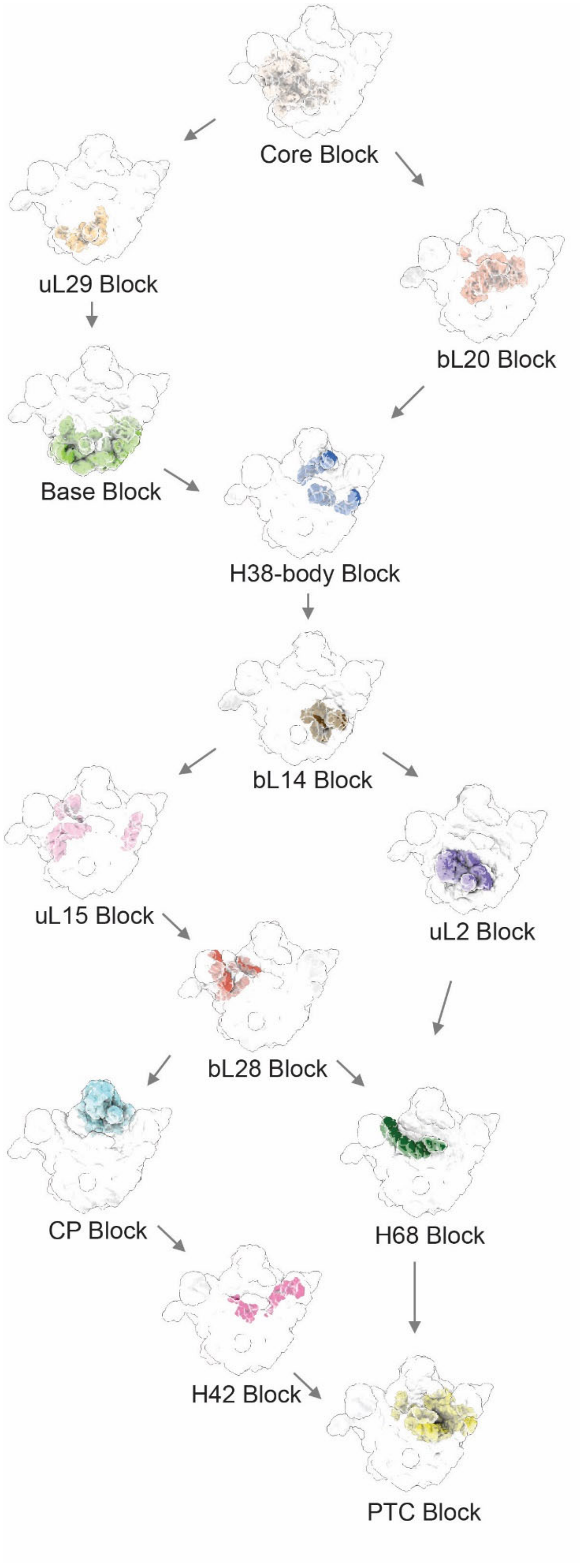

Core Block
uL29 Block
bL20 Block
Base Block
H38-body Block
bL14 Block
uL15 Block
uL2 Block
bL28 Block
CP Block
H68 Block
H42 Block
PTC Block

**Figure S4. Dependency map of iSAT assembly blocks.**

The dependencies of the assembly blocks are depicted in the dependency map, which is the same dependencies shown in Figure 4, showing electron density maps. The color-code is consistent with the previous figures.

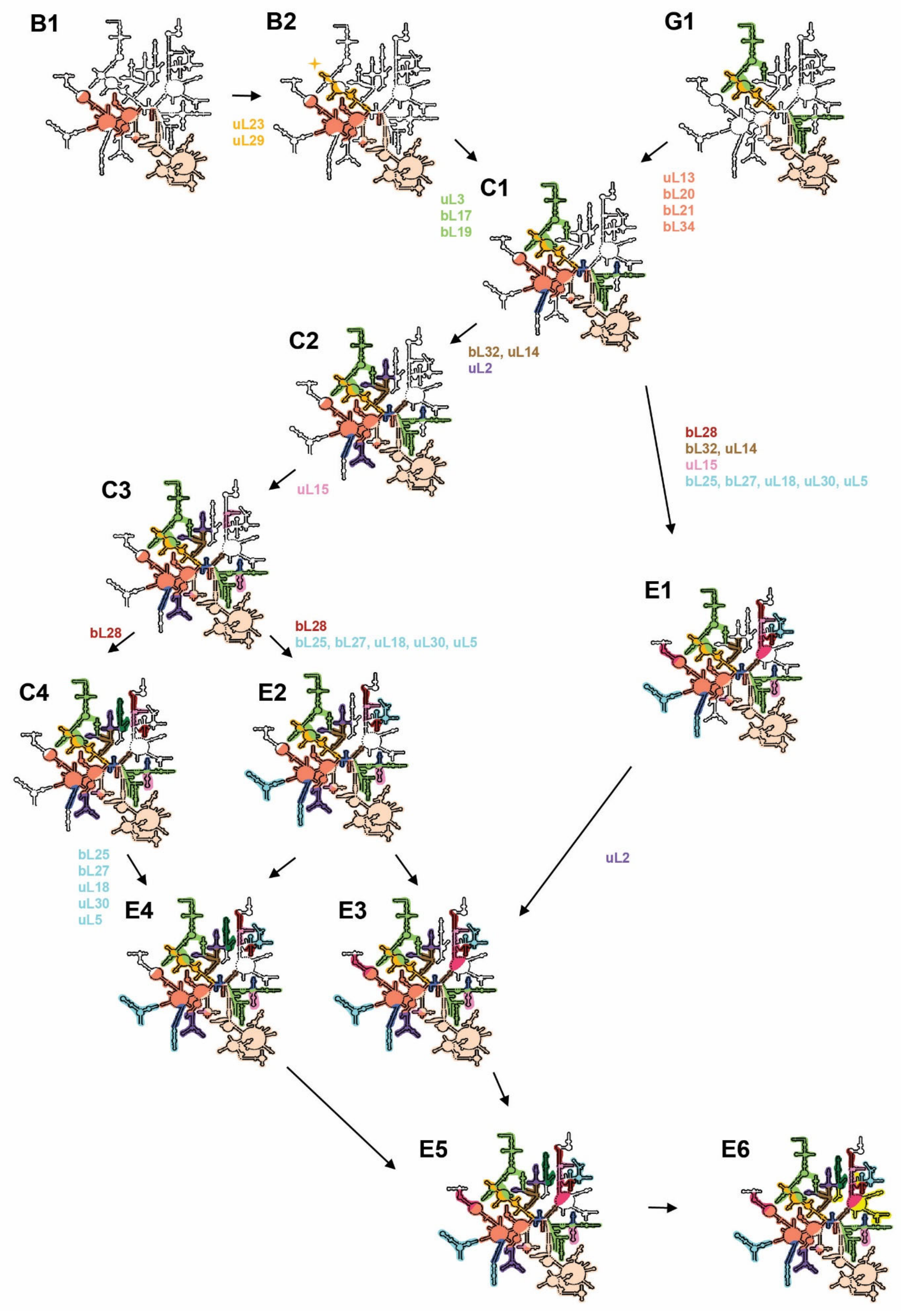
B1
B2
G1
uL23
uL29
C1
uL3
bL17
bL19
uL13
bL20
bL21
bL34
C2
bL32, uL14
uL2
bL28
bL32, uL14
uL15
bL25, bL27, uL18, uL30, uL5
C3
uL15
E1
bL28
bL28
bL25, bL27, uL18, uL30, uL5
C4
E2
bL25
bL27
uL18
uL30
uL5
uL2
E4
E3
E5
E6

**Figure S5. Assembly pathway highlighting RNA secondary structure elements.**

The thirteen reconstructed 50S **intermediates** from iSAT reaction time course are organized according to the dependencies of the assembly blocks, shown in Figure 7. The arrows indicate the minimal folding steps among the 50S precursors. Each 50S precursors are depicted on the rRNA secondary structure maps with colored correlated with assembly blocks in the previous figures. Star symbols highlight the new assembled helices between the two precursors. The new bound r-proteins during the transition of two precursors are also indicated near the arrows.

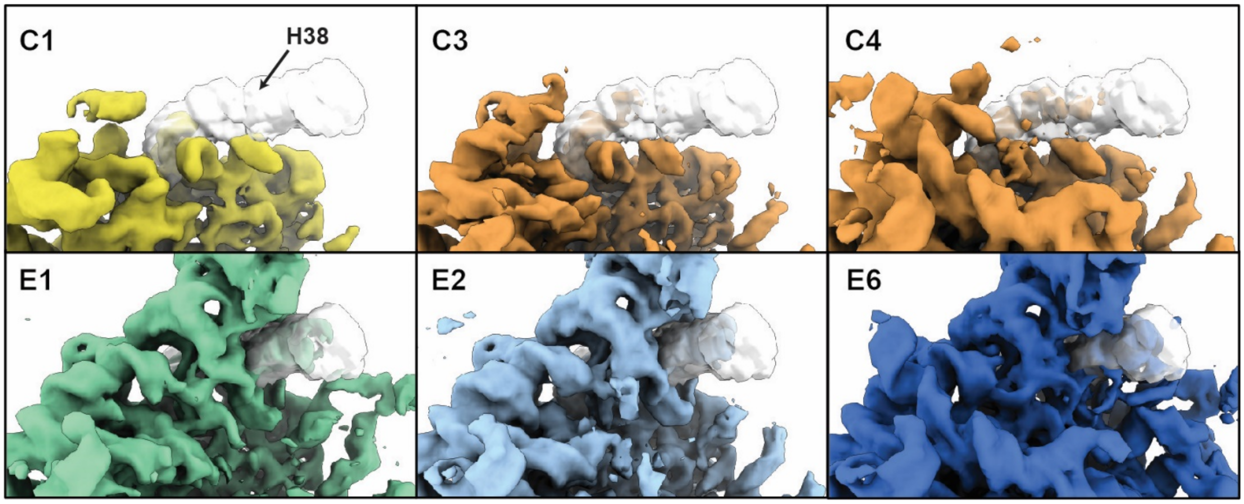


**Figure S6. Gradual organization of H38 during assembly.**

The transparent regions outlined in the panels are the electron density of Helix 38 in the crystal structure (PDB 4ybb). The colored densities are the density maps from iSAT time course dataset, with the precursors sorted in an immature-to-mature order.  There is a clear progression of maturity of Helix 38 with maturity of the class, which is relatively unique among the helical elements seen in this and other datasets.

### SUPPLEMENTARY TABLES

#### Table S1. rRNA modification inventory.

16S and 23S modifications identified and quantified using LC-MS relative to $^{15}$N-labeled 70S from E. coli MRE-600 cells. Sucrose gradient fractions, enriched in 70S ribosomes, were combined from two independent iSAT reaction preparations (**Replicate 1** and **Replicate 2**). Each sample has been treated with either RNase T1 or A. In **Replicate 2** sample, rRNA from iSAT and MRE 600 were combined and then treated with CMCT reagent before nuclease digestion. This enabled direct identification and quantification of pseudouridine modifications in addition to base and ribose methylation. Average **Fraction Modified** values over multiple charge states observed or multiple nuclease treatment (T1 and A) are reported when applicable. Empty cells indicate modifications not detected by LC-MS. Asterisk highlights values reported for a group of proximal modifications, that could not be resolved individually. For example, 40-50% of iSAT 23S molecules contain each of the following modifications: $m^1G$(745), Ψ(746), $m^5U$(747).

| | | | | Replicate 1 | | Replicate 2 | |
|---|---|---|---|---|---|---|---|
| | **position** | **modification** | **Enzyme** | **RNase used** | **Fraction modified** | **RNase used** | **Fraction modified** |
| | 516 | Ψ | RsuA | | | | |
| **Modifications of the 16S rRNA** | 527 | $m^7G$ | RsmG | T1 | 0.91 | | |
| | 966 | $m^2G$ | RsmC | T1 | 0.91 | T1 | 0.89 |
| | 967 | $m^5C$ | RsmD | T1 | 0.84 | T1 | 0.79 |
| | 1207 | $m^2G$ | RsmB | A | 0.86 | A | 0.88 |
| | 1402 | $m^4Cm$ | RsmH, RsmI | A | 0.48 | | |
| | 1407 | $m^5C$ | RsmF | T1 | 0.66 | | |
| | 1498 | $m^3U$ | RsmE | T1, A | 0.8 | T1, A | 0.88 |
| | 1516 | $m^2G$ | RsmJ | T1, A | 0.95 * | T1, A | 0.87 * |
| | 1518 | $m^6{}_2A$ | RsmA | T1, A | 0.95 * | T1, A | 0.87 * |
| | 1519 | $m^6{}_2A$ | RsmA | T1, A | 0.95 * | T1, A | 0.87 * |
| **Modifications of the 23S rRNA** | 745 | $m^1G$ | RlmA | T1 | 0.46 * | T1, A | 0.41 * |
| | 746 | Ψ | RluA | T1 | 0.46 * | T1, A | 0.41 * |
| | 747 | $m^5U$ | RlmC | T1 | 0.46 * | T1, A | 0.41 * |
| | 955 | Ψ | RluC | | | | |
| | 1618 | $m^6A$ | RlmF | T1 | 0.72 | | |
| | 1835 | $m^2G$ | RlmG | | | | |
| | 1911 | Ψ | RluD | T1, A | 0.54 * | T1 | 0.43 * |
| | 1915 | $m^3Ψ$ | RluD, RlmH | T1, A | 0.54 * | T1 | 0.43 * |
| | 1917 | Ψ | RluD | T1, A | 0.54 * | T1 | 0.43 * |
| | 1939 | $m^5U$ | RlmD | T1, A | 0.56 | T1 | 0.68 |
| | 1962 | $m^5C$ | RlmI | T1 | 0.9 | | |
| | 2030 | $m^6A$ | RlmJ | T1 | 0.66 | T1 | 0.98 |
| | 2069 | $m^7G$ | RlmKL | A | 0.56 | | |

| | | | | | | | |
|---|---|---|---|---|---|---|---|
| | 2251 | Gm | RlmB | | | | |
| | 2445 | $m^2G$ | RlmKL | | | | |
| | 2449 | hU | uknown | | | | |
| | 2457 | Ψ | RluE | | | A | 0.76 |
| | 2498 | Cm | RlmN | A | 1.17 | A | 0.89 |
| | 2501 | $ho^5C$ | RlhA | | | | |
| | 2503 | $m^2A$ | RlmN | A | 0.22 * | A | 0.33 * |
| | 2504 | Ψ | RluC | A | 0.22 * | A | 0.33 * |
| | 2552 | Um | RlmE | T1, A | 0.58 | A | 0.31 |
| | 2580 | Ψ | RluC | | | A | 0.9 |
| | 2604 | Ψ | RluF | | | T1 | 0.22 * |
| | 2605 | Ψ | RluB | | | T1 | 0.22 * |

**Table S2. rRNA and r-protein composition for assembly blocks.**

| Block Name | rRNA | r-protein |
|---|---|---|
| Core Block | H2-H24, H27-H28 | uL22, uL24, uL4 |
| uL29 Block | H47-H51, H53, H60, H104-H105 | uL23, uL29 |
| bL20 Block | H25-H26, H29, H32, H36-H37, H39-H41, H45-H46 | bL20, bL21, bL34, uL13 |
| Base Block | H1, H52, H54-H59, H63, H94, H96, H98-101, H106-H107 | bL17, bL19, uL3 |
| H95 Block | H38(body), H95, H102, H72 | |
| bL14 Block | H61, H64, H73 | bL32, uL14 |
| uL2 Block | H62, H65-H67, H33-H35, H103 | uL2 |
| uL15 Block | H31, H75, H79, H97 | uL15 |
| bL28 Block | H76, H88 | bL28 |
| H68 Block | H68-H70 | |
| CP Block | H38(cp), H108-H112, H81-H87 | bL25, bL27, uL18, uL30, uL5 |
| H42 Block | H74, H42 | |
| PTC Block | H71, H80, H89-H93 | bL35, bL36, uL16, uL6 |
| Stalks Block | H43-44, H77-78 | L1, L9, L10, L11, L7/12, L31 |

**Table S3. Particle numbers of thirteen intermediate classes from five time points**

| | 15min | 35min | 71min | 130min | 240min | Total |
|---|---|---|---|---|---|---|
| **B1** | 159 | 1162 | 966 | 620 | 148 | 3055 |
| **B2** | 491 | 3313 | 2470 | 1914 | 393 | 8581 |
| **G1** | 305 | 665 | 606 | 340 | 199 | 2115 |
| **C1** | 577 | 2159 | 1506 | 1075 | 488 | 5805 |
| **C2** | 2261 | 10120 | 10835 | 5967 | 2605 | 31788 |
| **C3** | 3283 | 2096 | 2069 | 1354 | 1105 | 9907 |
| **C4** | 184 | 389 | 467 | 282 | 306 | 1628 |
| **E1** | 577 | 2141 | 1010 | 285 | 201 | 4214 |
| **E2** | 341 | 814 | 1003 | 598 | 706 | 3462 |
| **E3** | 2216 | 2587 | 2552 | 1587 | 2551 | 11493 |
| **E4** | 367 | 631 | 986 | 488 | 604 | 3076 |
| **E5** | 709 | 1476 | 1730 | 1280 | 1791 | 6986 |
| **E6** | 536 | 2484 | 2377 | 1816 | 3540 | 10753 |
| **Total** | 12006 | 30037 | 28577 | 17606 | 14637 | 102863 |
